# White matter brain aging In Relationship to Schizophrenia and Its Cognitive Deficit

**DOI:** 10.1101/2020.10.19.344879

**Authors:** Jingtao Wang, Peter Kochunov, Hemalatha Sampath, Kathryn S. Hatch, Meghann C. Ryan, Fuzhong Xue, Jahanshad Neda, Thompson Paul, Britta Hahn, James Gold, James Waltz, L. Elliot Hong, Shuo Chen

## Abstract

We hypothesized that cerebral white matter deficits in schizophrenia (SZ) are driven in part by accelerated white matter aging and are associated with cognitive deficits. We used machine learning model to predict individual age from diffusion tensor imaging features and calculated the delta age (Δage) as the difference between predicted and chronological age. Through this approach, we translated multivariate white matter imaging features into an age-scaled metric and used it to test the temporal trends of accelerated aging-related white matter deficit in SZ and its association with the cognition. Followed feature selection, a machine learning model was trained with fractional anisotropy values in 34 of 43 tracts on a training set consisted of 107 healthy controls (HC). The brain age of 166 SZs and 107 HCs in the testing set were calculated using this model. Then, we examined the SZ-HC group effect on Δage and whether this effect was moderated by chronological age using the regression spline model. The results showed that Δage was significantly elevated in the age >30 group in patients *(p* < 0.001) but not in age ⩽ 30 group *(p* = 0.364). Δage in patients was significantly and negatively associated with both working memory *(β* = −0.176, *p* = 0.007) and processing speed *(β* = −0.519, *p* = 0.035) while adjusting sex and chronological age. Overall, these findings indicate that the Δage is elevated in SZs and become significantly from middle life stage; the increase of Δage in SZs is associated with the decline neurocognitive performance.

## 1. Introduction

Patients with schizophrenia (SZ) are at risk for elevated aging-related functional and neurological decline, termed accelerated aging (Kirkpatrick et al., 2008; Ito and Barnes, 2009; Jeste et al., 2011; Kochunov et al., 2013b, 2014, 2016b). They are at significantly higher risks (1.5 to 5 times) for developing cognitive deficits and dementia before age 70 (Cai and Huang, 2018; Chen et al., 2015; Diniz et al., 2017; Ribe et al., 2015). Previous studies demonstrated that the integrity of the cerebral white matter as measured by fractional anisotropy (FA) in diffusion weighted imaging (DTI), declined in SZ patients at nearly twice the aging rate of normal controls (Mori et al., 2007; Friedman et al., 2008; Kochunov et al., 2013a, 2013b; Wright et al., 2014). An analysis of life-long trajectory of white matter integrity in 600 schizophrenia patients suggested that the peak of white matter integrity occurs earlier in SZ than healthy controls and the accelerated decline of the associative white matter tracts becomes evident in the 4^th^ decade of life and its slope shows a non-linear increase with age (Cetin-Karayumak et al., 2019).

The strong aging related sensitivity of white matter measures such as FA (Kochunov et al., 2016a) can also be used to predict “brain age” for individual subjects using neuroimaging data. The difference between brain age and chronological age can then be used as a phenotype to evaluate evidence for accelerated or slower aging in an individual (Cole and Franke, 2017; Franke et al., 2012; Smith et al., 2019; Wang et al., 2019). The “brain age” analysis can be performed using machine learning and/or regression models that are trained to draw association between regional brain measures and chronological age. The delta age (Δage), the difference between brain age and chronological age, is expected to be null on average in a group of individuals undergoing the normal brain aging process. It translates multivariate imaging features into an age-scaled metric that can be used as an index to depict imaging-based brain structural changes during aging. The Δage may be increased by the atypical brain aging caused by physical and brain diseases (Franke et al., 2013, 2012) such as dementia (Wang et al., 2019), Alzheimer’s disease (Gaser et al., 2013), schizophrenia (Koutsouleris et al., 2014; Nenadić et al., 2017), or epilepsy (Holmes et al., 2012).

Patients with schizophrenia (SZ) have a significantly higher risk (2-4 times) of experiencing an unfavorable aging trajectory that may increase their risk for cognitive decline and dementia (Ribe et al., 2015). The increased risk can be in part explained by the fact that the white matter integrity reaches its peak earlier in life for SZ than healthy controls (Cetin-Karayumak et al., 2019). Neuropathology of accelerated brain aging likely contributes to the severity of the cognitive deficits in patients that form the core of socioeconomic impairments in this illness (Kochunov et al., 2016b; Kelly et al., 2018; Kochunov et al., 2017). Previously, we have shown that white matter deficits are associated with cognitive deficits and treatment difficulty in SZ (Kochunov et al., 2019, 2017, 2016b) and have hypothesized that accelerated white matter aging in SZ leads to development of cognitive deficits and disorder-specific deficit patterns (Kochunov and Hong, 2014).

Built on the previous research, we used machine learning to calculate white matter Δage and further tested two hypotheses that (1) white matter Δage is elevated in patients with schizophrenia and the SZ-related elevation in Δage is moderated by chronological age (i.e., white matter Δage in SZ across the lifespan); and (2) the white matter Δage is associated with the core cognitive deficits in schizophrenia while adjusting chronological age and other covariates.

## 2. Materials and methods

### 2.1. Participants

All the patients were recruited in Maryland Psychiatric Research Center, University of Maryland School of Medicine and neighboring outpatient clinics. Healthy controls were recruited via advertising, e.g., flyers, social media and word of mouth. Imaging data were available for N = 166 individuals with diagnosis of schizophrenia spectrum disorders (117 Male / 49 Female, average age = 36.232) and 214 HC (140 Male / 74 Female, average age = 38.016) without current Axis I psychiatric illnesses (Table 1). Among all SZs, 11 subjects were diagnosed as schizoaffective disorder while others were diagnosed as schizophrenia. Diagnosis of schizophrenia spectrum disorders was conducted according to Diagnostic and Statistical Manual of Mental Disorders-IV (DSM-IV) criteria through a best estimate approach combining information from a Structured Clinical Interview for DSM-IV (SCID) with a review of medical records. The exclusion criteria included diagnosis with hypertension, hyperlipidemia, type 2 diabetes, heart disorders, major neurologic event such as stroke or transient ischemic attack, and recent substance use disorder (except tobacco and marijuana use). It was applied for all participants based on self-reported questionnaires. All participants gave written informed consent approved by the University of Maryland Baltimore institutional review board.

**Table 1.**
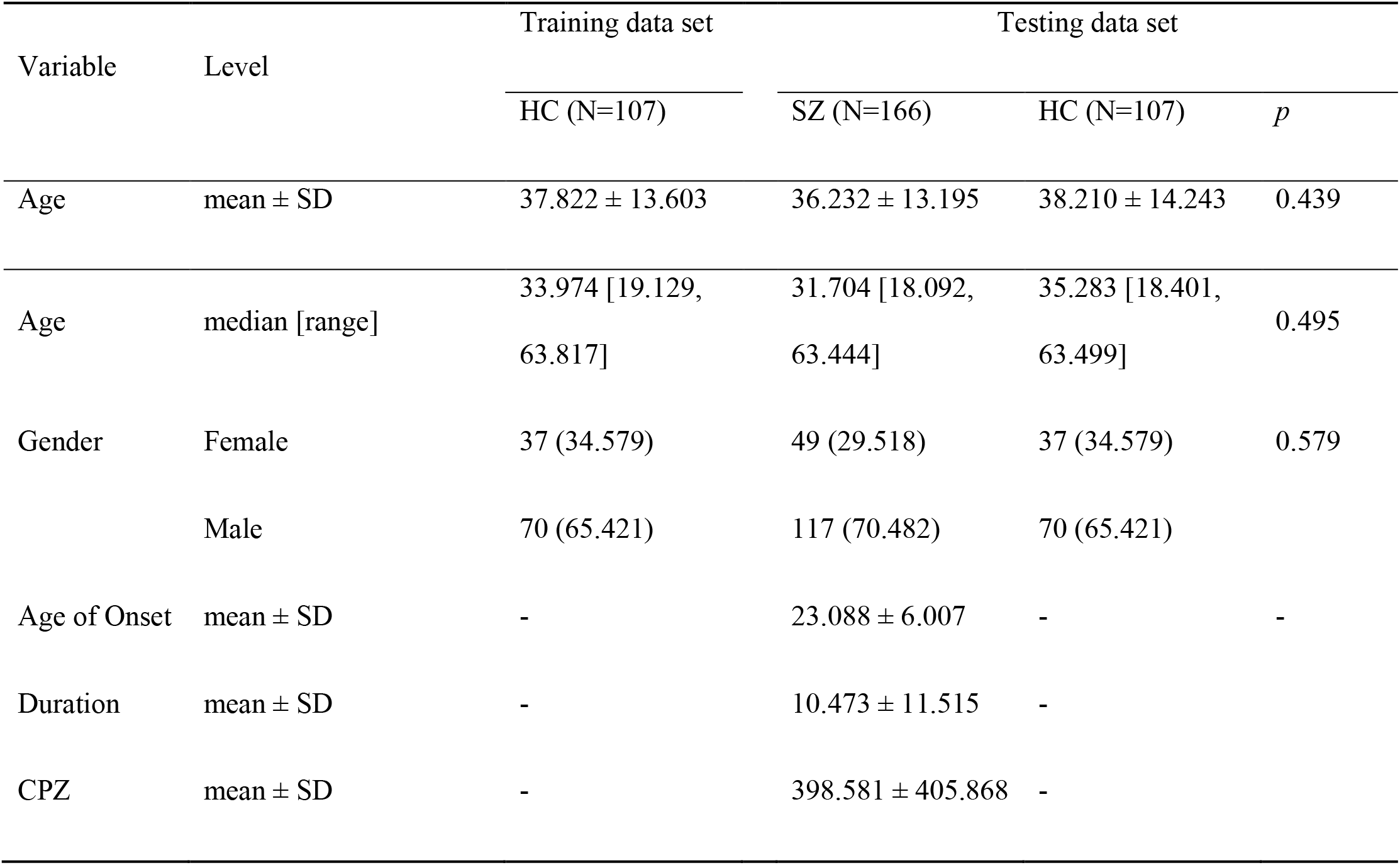
Demographics and clinical characteristics of participants

To evaluate the prediction performance of the machine learning model, two parts were extracted from HC frequency-matching the distribution of SZ subjects’ age and gender. One part consisted of 107 HC (70 Male / 37 Female, average age = 37.822) was used to fit the machine learning model. The other part consisted of 107 HC (70 Male / 37 Female, average age = 38.210) constituted the testing set with SZ subjects together. Additional clinical and epidemiologic information is provided in Table 1.

### 2.2. Image Acquisition and Processing

Magnetic resonance images were acquired through studies using magnetic resonance imaging 3T Siemens scanner. Imaging data was collected using a Siemens 3T TRIO MRI (Erlangen, Germany), running an upgraded VB17 software and a 32-channel RF head coil. DTI data was collected using a spin-echo, EPI sequence with a spatial resolution of 1.7×1.7×3.0 mm. The sequence parameters were: TE/TR = 87/8000 ms, FOV = 200 mm, axial slice orientation with 50 slices and no gaps, 64 isotropically distributed diffusion weighted directions, two diffusion weighting values (b = 0 and 700 s/mm2) and five b = 0 images. Subjects’ head movement was minimized with restraining padding. The DTI data were processed using the ENIGMA DTI analysis pipeline (https://www.nitrc.org/projects/enigma_dti) (Jahanshad et al., 2013). All data included in the analysis passed the ENIGMADTI quality assurance or quality control procedures. Regional white matter FA measurements were generated for 43 tracts (Supplementary methods). Among them, the per-tract mean values were found by calculating the mean values along tract regions of interest per hemisphere except for commissural tracts.

### 2.3. Data Analysis and Statistical Modeling

#### 2.3.1 Machine learning model for white matter brain age calculation

A machine learning (ML) i.e. supervised learning computational model is first used to calculate ‘white matter brain age’ of each participant using only DTI features. This task can be implemented by training an optimal machine learning model in community control participants (Cole and Franke, 2017; Liem et al., 2017; Shahab et al., 2019), locking the model, and applying the model to patients and controls in the testing sample. The training of the ML model is often based on solely the controls (Franke et al., 2012; Khundrakpam et al., 2015), same as we have done here (N = 107). In the training stage, chronological age was considered as the continuous outcomes/labels (Y) and white matter FA values as the input features (X). To achieve the best prediction performance, several machine learning model candidates including Random forest regression (Breiman, 2001; Geurts et al., 2006), Gradient boosting regression(Friedman, 2002, 2001), LASSO (Least absolute shrinkage and selection operator) (Friedman et al., 2010) and others were utilized and compared. The feature selection, parameter tuning of each model and model comparison were conducted using 5-fold cross-validation (CV) within the training data set. The criteria of prediction performance were the coefficients of determination (*R*^2^) between the chronological and brain age (which is equivalent to the mean absolute error (MAE) criteria) (Fig 1A). The best performing model in the CV then was locked and applied to the testing sample, thus the testing sample results were not influenced by the training data. Note that the demographic variables are balanced between the training and testing data sets, and between testing HC and SZ groups.

**Fig. 1.**
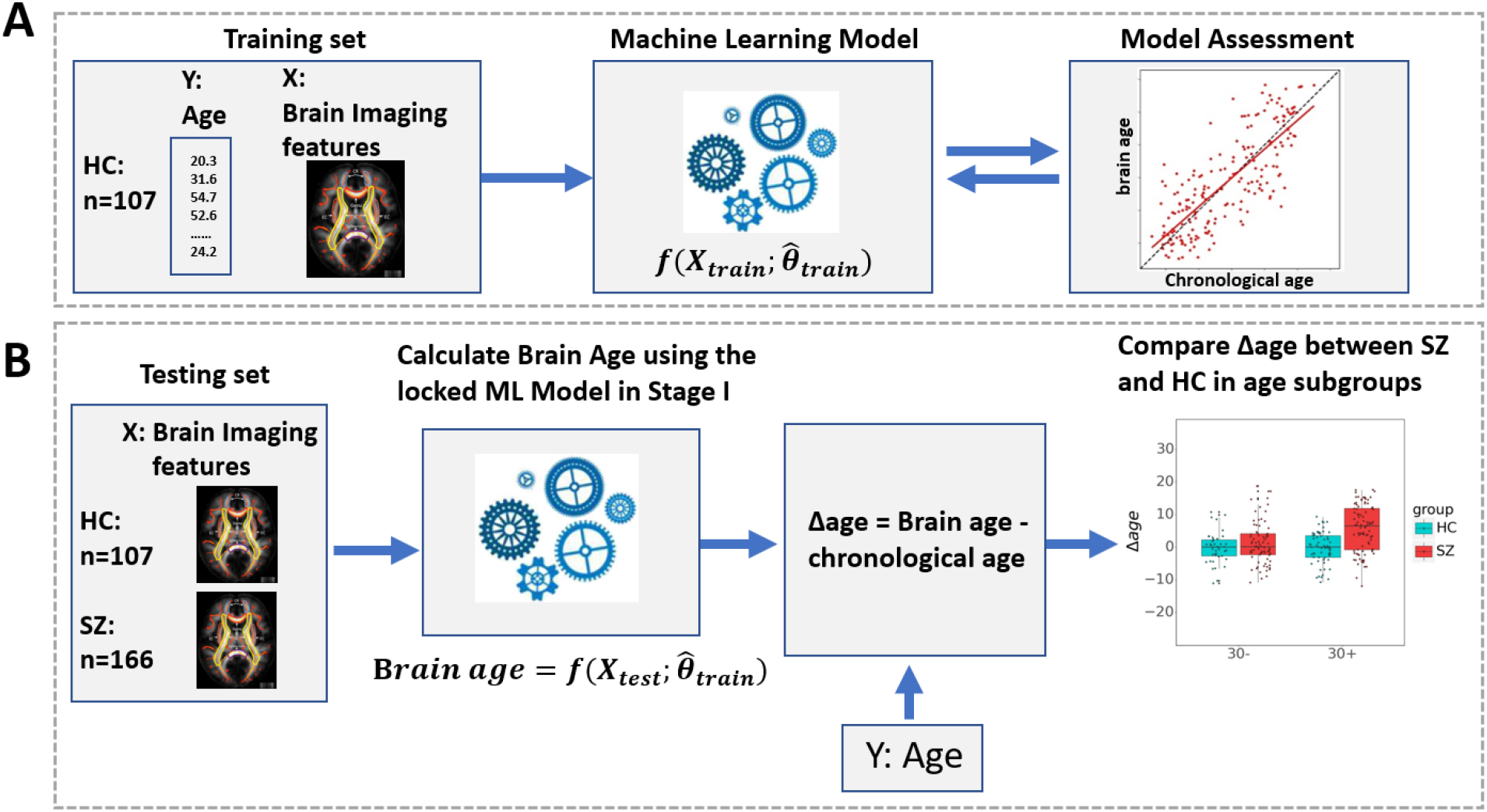
Flowchart showing the process of study design. (A) Training: Machine learning model estimation for brain age. The optimal machine learning model in 5-fold cross-validation was locked. The parameters of the optimal machine learning model were designated as 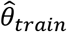. (B) Testing: Compare brain age between HCs and SZs in different age groups. The FA values of HCs and SZs in the testing data set were subjected to the locked random forest model to predict the brain age. The Δage was compared between HCs and SZs in the testing data set in two age groups.

Next, the locked machine learning model was applied to the testing data set which includes both SZ (N = 166) and HC (N = 107). The model was used to calculate the ‘brain age’ based on the input brain imaging features (Fig 1B). The brain age delta (Δage) was then calculated. A positive value would suggest an accelerated white matter aging effect.

#### 2.3.2 Statistical analysis of Δage

We first tested whether white matter Δage is associated with SZ, and whether the diagnosis group effect is moderated by chronological age. Since the moderation effect of chronological age is not linear, we applied a comprehensive and data-driven approach to objectively determine the disease effect on Δage across the lifespan. Specifically, we used spline models including linear splines and natural cubic splines as basis functions to assess the moderation effect of the chronological age (Wood, 2011). Then, we performed model selection and chose the optimal location and number of knots and spline basis function based on the likelihood criteria (e.g., BIC). The final model can provide statistical inference to evaluate the effect of SZ on Δage across the lifespan. We then tested the difference of Δage between SZ and HC in each age subgroup using two sample tests after correction for multiple comparisons; note that age and sex were frequency-matched between SZ and HC.

We also examined the association between Δage and cognitive deficits in schizophrenia patients. The Wechsler Abbreviated Scale of Intelligence digit-sequencing and digit-symbol-coding subscales were used to assessed working memory and processing speed, respectively (Wechsler, 1999); these tasks were selected as they are among the most robust tasks separating those with schizophrenia spectrum disorders vs. controls in meta-analysis across all cognitive domains (Dickinson et al., 2007). These metrics were regressed against Δage independently using the general linear model (GLM). Since Δage was hypothesized to be elevated in SZ but was expected to be null on average in HC along lifespan, we assessed the impact of Δage on cognitive deficits in both SZ and HC group (Wang et al., 2007). Chronological age and sex were adjusted as covariates. In addition, to assess the robustness of results, we repeated the statistical analysis by excluding the schizoaffective disorder patients.

## 3. Results

### 3.1. Participants characteristics

The demographic characteristics are shown in Table 1. No significant difference was found in age and gender between the three parts of data set *(p* = 0.439, *p* = 0.579, respectively).

### 3.2. Machine learning model selection on the training data set

The random forest regression model achieved the best performance in predicting the age in the test-retest trials in HC participants. The features selection in random forest model was conducted using the recursive feature elimination (Breiman, 2001; Geurts et al., 2006). Among 43 tracts, FA values of 34 tracts were selected (Supplementary methods). The parameter tuning of random forest was conducted using 5-fold CV with the *R*^2^ as the criteria of prediction performance. The random forest model predicted chronological age with an *R*^2^ of 0.916 (MAE = 4.640 years, *p* < 0.001) and achieved good performance when be applied to HC (*R*^2^ = 0.895, MAE = 3.649 years, *p* < 0.001) and SZ (*R*^2^ = 0.814, MAE = 6.469 years, *p* < 0.001) in the testing dataset (Supplementary Fig. S1). The performance of other models was inferior and presented in the supplementary Table S1.

### 3.3. Accelerated aging in patients

The correlation between chronological age and Δage were significantly in SZ (*R*^2^ = 0.048, *p* = 0.005) but not significantly in HC (*R*^2^ = 0.010, *p* = 0.296) in the testing dataset (Supplementary Fig. S2). The effect of SZ by age interaction on Δage was positive and significant (*β* = 0.134, *p* = 0.007), which indicated the effect of group on Δage is moderated by chronological age. Furthermore, we examined the SZ-HC group effect on Δage, and the nonlinear moderation effect of chronological age on the group effect using the regression spline model. Based on the likelihood criteria, the final model which fitted the group effect on Δage across the lifespan optimally is a linear spline model with one knot at age = 30. Based on this age knot, the chronological age was divided into two subgroups (≤30 years old and > 30 years old). The Δage was then compared between SZs and HCs for these two age groups. The Δage of the SZs was not significantly higher than that of HCs in ≤30 age group (mean difference (95% CI): 1.479 (−0.586, 3.543), *p* = 0.364) but was significantly higher in >30 age group (5.903 (3.989, 7.817), *p* < 0.001) (Fig. 2). The demographic characteristics of testing set in two age groups are presented in Table 2.

**Fig. 2.**
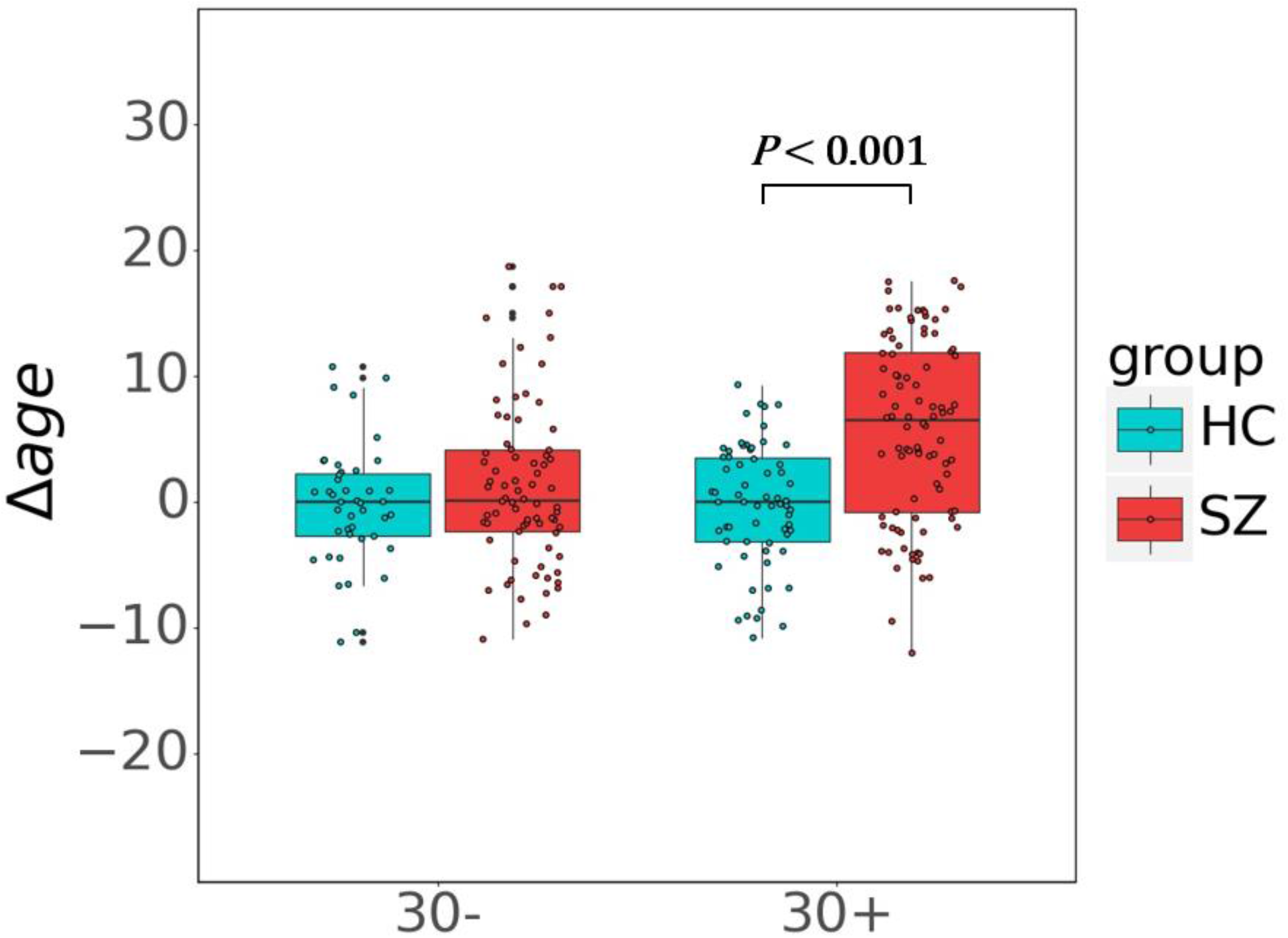
Box plots demonstrate the differences of Δage between SZ and HC in the testing data set for two age groups. Significant Δage difference between SZ and HC emerged at age group 30+. No significant differences were found in the younger age group.

**Table 2:** Demographics and clinical characteristics of participants in each age group in testing set

| Variable | Level | age ≤ 30 | | age > 30 | |
| --- | --- | --- | --- | --- | --- |
|  |  | SZ (N=76) | HC (N=43) | SZ (N=90) | HC (N=64) |
| Age | mean ± SD | 24.204 ± 3.374 | 23.74 ± 3.145 | 46.389 ± 9.238 | 47.932 ± 9.788 |
|  | median | 24.726 [18.092, | 23.606 [18.401, | 48.168 [30.313, | 48.905 [30.226, |
|  | [range] | 29.610] | 29.788] | 63.444] | 63.499] |
| Gender | Female | 18 (23.684) | 15 (34.884) | 31 (34.444) | 22 (34.375) |
|  | Male | 58 (76.316) | 28 (65.116) | 59 (65.556) | 42 (65.625) |
| Age of Onset | mean ± SD | 18.536 ± 3.024 | - | 22.913 ± 8.005 | - |
| CPZ | mean ± SD | 340.169 ± 340.916 | - | 554.702 ± 443.104 | - |

### 3.4. Assessing the correlation between Δage and cognition

In patient, Δage was significantly associated with both working memory *(β* = −0.176, *p* = 0.007) and processing speed *(β* = −0.519, *p* = 0.035) after adjusting sex and chronological age (Table 3). Average daily antipsychotic medications as measured by Chlorpromazine equivalent (CPZ) was not significantly associated with Δage in the patients *(β* = 0.002, *p* = 0.179). Age of psychosis onset in the patients was also not significantly associated with Δage *(β* = −0.007, *p* = 0.970). Finally, duration of the illness in the patients was also not significantly associated with Δage *(β* = 0.110, *p* = 0.165). In the controls, neither working memory *(β* = −0.059, *p* = 0.636) nor processing speed measures *(β* = −0.180, *p* = 0.618) was significantly associated with Δage after adjusting sex and chronological age (Table 3, Fig 3). The results showed that Δage and cognitive deficits are correlated only for patients with schizophrenia, but not for healthy controls. We also performed regression analysis using the combined SZ and HC cohort (N=273) with covariates of chronological age, Δ age, sex, group, chronological age × group, and Δ age × group. The results were similar to the subgroup analysis and summarized in Supplementary table S2. In addition, we repeated the statistical analysis by excluding the schizoaffective disorder patients, and the results remained similar to the main results (supplementary materials).

**Fig. 3.**
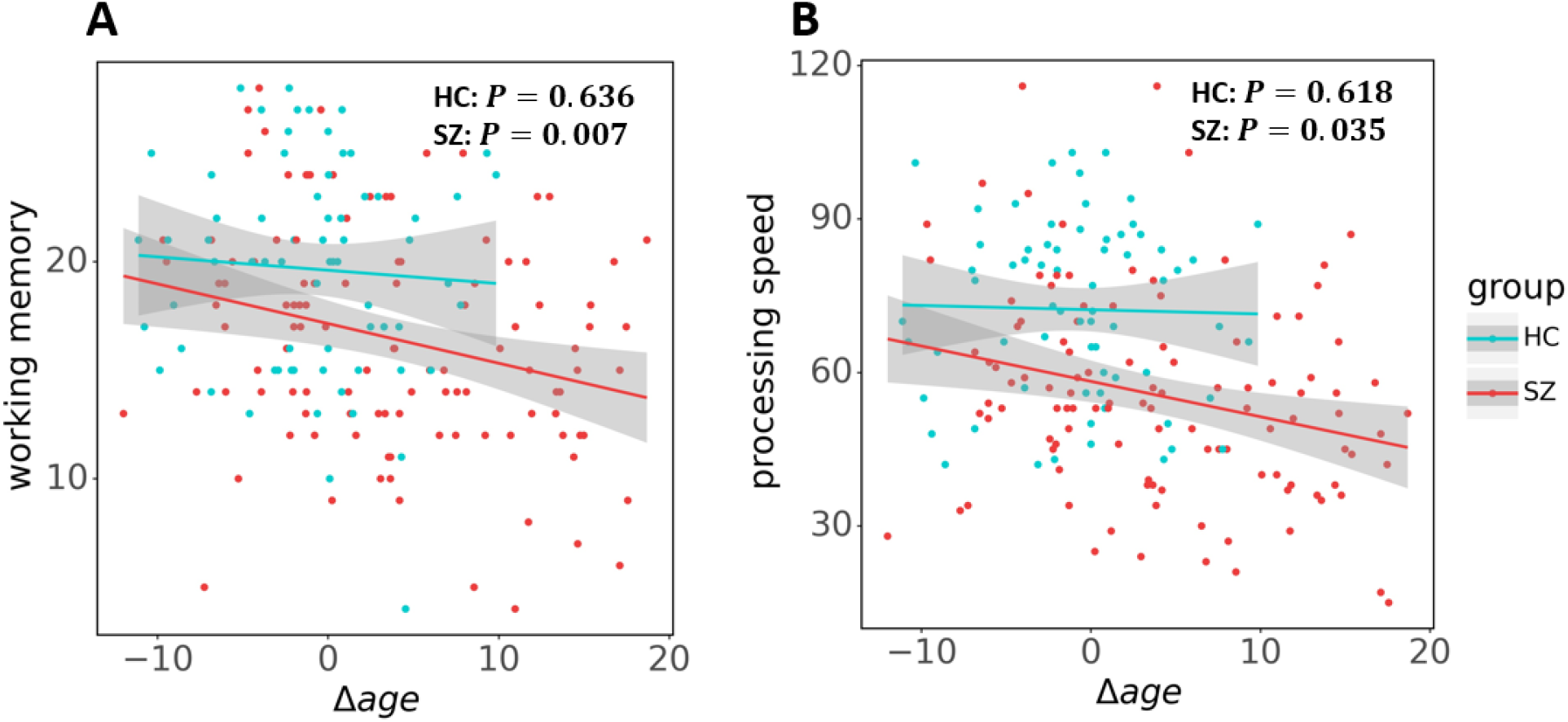
Scatterplots and linear fits illustrating relationships between Δage and working memory and processing speed in patients with schizophrenia and healthy controls.

**Table 3.** Associations between two age-scaled metrics and cognition

| Group | Variable | Working memory |  | Processing speed |  |
| --- | --- | --- | --- | --- | --- |
| | | $\beta$ | $p$ | $\beta$ | $p$ |
| SZ | Sex | -1.530 | 0.147 | 7.784 | 0.053 |
|  | chronological age | -0.067 | 0.053 | -0.350 | 0.009** |
| | Δage | -0.176 | 0.007** | -0.519 | 0.035* |
| HC | Sex | -0.900 | 0.502 | 1.999 | 0.604 |
|  | chronological age | -0.020 | 0.636 | -0.674 | <0.001*** |
| | Δage | -0.059 | 0.636 | -0.180 | 0.618 |

\* $p < .05$ , \*\* $p < .01$ , \*\*\* $p < .001$ .

## 4. Discussion

In the current study, a novel Brain Age analysis approach was applied to perform personalized predictions of individual age based on white matter imaging. The calculated white matter Δage, the difference between predicted and chronological ages, was elevated in SZ compared to healthy controls and the difference of Δage between SZ and HC was moderated by chronological age. Followed determining the optimal age knot based on the likelihood principle, we assigned participants into two age groups and found that Δage was significantly higher in SZ relative to HC at age group old than 30 years old but not significantly at age group younger than 30 years old. Furthermore, we also identified that Δage in patients was associated with worsened performance on two of the neurocognitive tests most affected in schizophrenia: working memory and processing speed.

Most recently, using machine learning to predict the age of an individual based on neuroimaging has received increased interest. An increasing number of studies tried to construct brain age in this approach based on varieties of neuroimaging data, such as diffusion tensor imaging, cortical thickness or cognitive performance scores, and further investigated whether the brain age of patients with schizophrenia were higher than those of the healthy controls. Three previous studies in SZ used Brain Age approach with structural T1w data and showed that the average Δage in patients was significantly higher (2.6 to 8.8 years) compared to healthy controls (Franke et al., 2010; Hajek et al., 2019; Steffener et al., 2016). Koutsouleris and colleagues also showed that patients with schizophrenia had significantly higher average Δage than patients with major depressive or borderline personality disorders (Koutsouleris et al., 2014). Hajek et al. reported a significantly higher Δage even in the first-episode schizophrenia patients compared with controls. Together, these studies provided strong evidence that participants with schizophrenia experience an accelerated aging compared to HCs, however the timing of these changes remained unclear (Hajek et al., 2019).

This is the first brain age analysis in SZ that was focused on cerebral white matter. We demonstrated that Δage was significantly greater in patients than healthy controls thus supporting previous reports of accelerated white matter aging in this illness (Cetin-Karayumak et al., 2019; Kochunov et al., 2013b). Moreover, we found no significant differences in brain age among young patients and Δage only became significant in middle age in SZ. Our age group analysis of Δage differences among two subgroups further suggests that accelerated decline in FA may represent the life-long interaction with schizophrenia. The lack of significant differences in Δage in ages ≤30 is interesting. A study by Cetin-Karayumak and colleagues summarized three possible trajectories reported in previous studies and investigated which white matter trajectory fitted best based on a large cohort. These three possible trajectories included: (1) “neurodevelopmental models in schizophrenia” postulating that schizophrenia is caused by environmental and/or genetic insults that occur during prenatal, perinatal, or early childhood/adolescence, leading to lower and parallel trajectories throughout the lifespan as compared to healthy controls; (2) “maturational model in schizophrenia” suggesting that disturbances during maturation would cause different ascending slopes and a shift in peak while the similar starting and end points of trajectories compared to healthy controls, indicating perturbed myelination; (3) “accelerated aging in schizophrenia” supporting that schizophrenia is marked by similar trajectory during maturation of white matter but steeper descending slopes related to accelerated aging processes, such as myelin breakdown. They found that whole-brain FA showed a monotonic increase until reaching a peak at the age of 33 years in controls, and earlier, at the age of 27 years in patients and the confidence interval of FA trajectory in two groups overlapped in this period. FA declined monotonically after reaching peak in both patients and controls, and demonstrated accelerated decline rate in patients compared to controls (Cetin-Karayumak et al., 2019). This finding supported the third possible trajectory, which may explain the similarity in Δage for younger age before the peak for cerebral white matter FA increases in patients and the accelerated increase in Δage of patients after 30 years old in our study.

Previous studies have reported that FA deficits are associated with cognitive deficits in schizophrenia (Epstein et al., 2014; Miyata et al., 2010; Nazeri et al., 2013; Pérez-Iglesias et al., 2010; Roalf et al., 2013; Voineskos et al., 2013). Karlsgodt and colleagues observed that lower FA values were predictive of negative changes in cognitive status (Karlsgodt et al., 2009). Previously, we have hypothesized that accelerated white matter aging in SZ may leads to development of cognitive deficits (Kochunov and Hong, 2014). In the current study, we calculated Δage through the novel Brain age analysis approach and tested its association with cognitive deficits, and showed that Δage in patients was significantly and negatively correlated with performance on the working memory and processing speed neurocognitive tasks. All correlations were negative, indicating that patients who were predicted to be older than their chronological age had worse neurocognitive performance on average. The neurocognitive deficits in these two domains of schizophrenia are enduring, pervasive, and form the core of the functional disability in patients (Dickinson et al., 2007; Faraone et al., 2000; Keefe et al., 2005, 2004; Knowles et al., 2010). The neurocognitive deficits in these domains have already been linked with reduced white matter integrity in patients, suggesting that integrity of long-distance neuronal fibers is critical to maintaining normal performance of long-distance cortical networks that serve these functions (Kochunov et al., 2017). Δage does not measure the underlying neurobiological mechanisms. However, it provides an aggregate measure of the white matter deficit pattern along chronological age. In our recent study, we observed the white matter deficit patterns in SZ are strikingly similar with that in Alzheimer disease and are associated with working memory and processing speed. Moreover, the similarity between the white matter deficit patterns in these two diseases were positively correlated with age (Kochunov et al., 2020). This increasing similarity along age may be reflected by Δage and hence cause the association between Δage and cognitive performance. The highest correlation was observed with the working memory function. Working memory deficits are among the core cognitive deficits reported in schizophrenia (Dickinson et al., 2007; Knowles et al., 2010). Working memory deficits are also directly linked to white matter in both subjects with schizophrenia and normal controls (Karlsgodt et al., 2009; Nazeri et al., 2013; Kochunov et al., 2017; Zeng et al., 2016; Karlsgodt et al., 2010).

There are several limitations in the current study. First, the cross-sectional design may limit our ability to fully rule out confounding effects from antipsychotic medications and differences in illness duration and age of onset on Δage in the patients, although they were not significantly associated with Δage in the current analysis. A longitudinal DTI study will be necessary to determine the white matter decline trajectories of normal individuals, which will enable estimation of more precisely the effect of schizophrenia on accelerated white matter aging and provide causal inference. Second, the etiology causing the pattern of accelerated white matter aging described here is unknown and not studied in the current study, although we found that current dose of antipsychotic medications that patients were on did not significantly contribute to their Δage.

In conclusion, we confirmed white matter based brain age in participants with schizophrenia was elevated compared with healthy controls in relationship to their chronological age and identified that this abnormality occurred at middle and older age and was not significant in young SZ patients. The white matter Δage can be used as a simple, individual-level indicator that reflects accelerated white matter brain aging in SZ based imaging features. The pattern of accelerated brain aging in SZ along the lifespan in our analysis can help to reveal the dynamic disease progression of SZ on brain aging and the associated reduction of neurocognitive performance.

