## Supplementary Materials for "White matter brain aging In Relationship to Schizophrenia and Its Cognitive Deficit"

### Supplementary Methods

#### Participants enrollment

In this analysis we used 81 HC overlapped with the previous publication (Kochunov et.al, JAMA Psychiatry, 2017) in our training data set, and 77 overlapped HC and 125 SZ in the testing data set.

#### White matter tracts used in the analysis

In this analysis regional white matter FA measurements were generated for 43 tracts including left anterior corona radiata (ACRL), right anterior corona radiata (ACRR), left anterior limb of internal capsule (ALICL), right anterior limb of internal capsule (ALICR), body of corpus callosum (BCC), corpus callosum (CC), left Cingulum (CGCL), right Cingulum (CGCR), left perihippocampal cingulum tract (CGHL), right perihippocampal cingulum tract (CGHR), left corona radiata (CRL), right corona radiata (CRR), left corticospinal tract (CSTL), right corticospinal tract (CSTR), left external capsule (ECL), right external capsule (ECR), fornix (FX), left fornix stria terminalis (FXSTL), right fornix stria terminalis (FXSTR), genu of corpus callosum (GCC), left internal capsule (ICL), right internal capsule (ICR), left inferior fronto occipital fasciculus (IFOL), right inferior fronto occipital fasciculus (IFOR), left posterior corona radiata (PCRL), right posterior corona radiata (PCRR), left posterior limb of internal capsule (PLICL), right posterior limb of internal capsule (PLICR), left posterior thalamic radiation (PTRL), right posterior thalamic radiation (PTRR), left retrolenticular part of internal capsule (RLICL), right retrolenticular part of internal capsule (RLICR), splenium of corpus callosum (SCC), left superior corona radiata (SCRL), right superior corona radiata (SCRR), left superior fronto-occipital fasciculus (SFOL), right superior fronto-occipital fasciculus (SFOR), left superior longitudinal fasciculus (SLFL), right superior longitudinal fasciculus (SLFR), left sagittal stratum (SSL), right sagittal stratum (SSR), left uncinate fasciculus (UNCL), right uncinate fasciculus (UNCR).

Followed features selection conducted using the recursive feature elimination with random forest model, FA values of 34 tracts including ACRL, ACRR, CC, CGCL, CGCR, CGHL, CRL, CRR, CSTL, CSTR, ECL, FX, FXSTL, FXSTR, GCC, ICR, IFOL, IFOR, PCRL, PCRR, PLICR, PTRL, PTRR, RLICL, SCC, SCRL, SCRR, SFOL, SFOR, SLFL, SLFR, SSL, SSR, UNCL were used to fit the brain age prediction model..

### Supplementary Figures

#### Supplementary Figures S1

*Predicted Brain age*

As shown in Figure S1, the linear model of brain-predicted age with chronological age in HC and SZ. The overall coefficient between brain-predicted age and chronological age in SZ and HC are 1.120 and 0.966, respectively. The coefficients of two linear model was tested using *F*-test ($F(2, 269) = 16.715$, $P < 0.0001$). In young subjects, the brain-predicted ages of SZ didn't significantly differ with HCs. The brain-predicted age of schizophrenia patients begins to be significantly greater than HC test from 30 years old. As chronological age increases, the gap of brain-predicted age between SZ and HC are larger and larger.

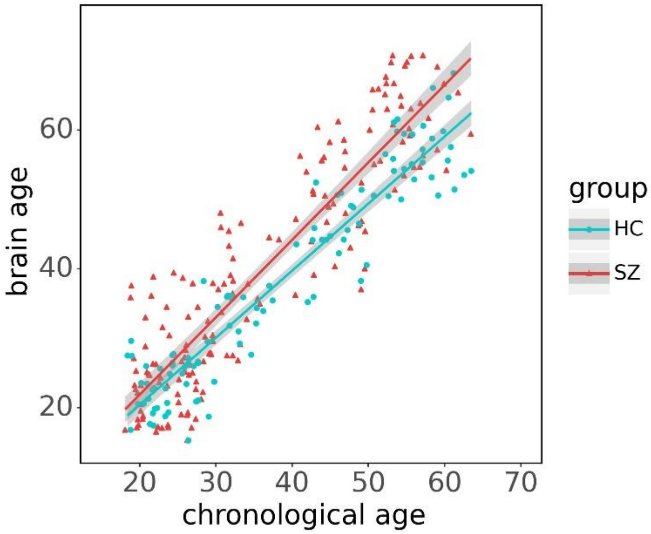


Fig. S1. Brain-predicted age and chronological age in HC and SZ with the confidence interval.

*△age in SZ and HC across lifespan*

As shown in Figure S2, the linear model of △age with chronological age in HC and SZ. The overall coefficient between brain-predicted age and chronological age in SZ and HC are 0.120 and -0.034 respectively. The coefficients of two linear model was tested using *F*-test ($F(2, 269) = 16.715$, $P < 0.0001$). As chronological age increases, the gap of brain-predicted age between SZ and HC are larger and larger.

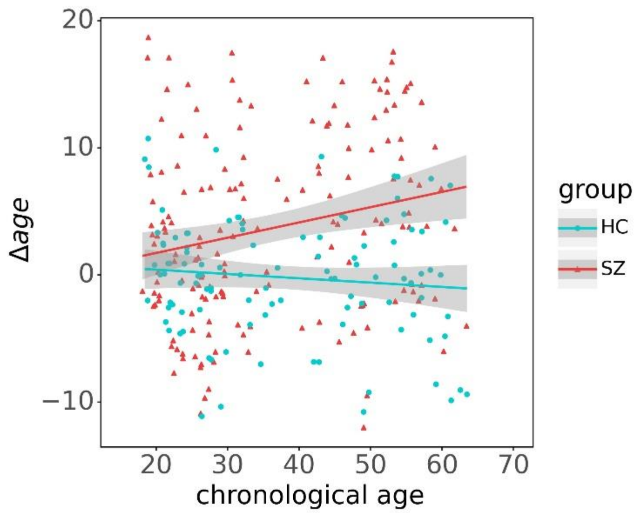


Fig. S2. Δage and chronological age in HC and SZ with the confidence interval.

### Supplementary Tables

#### Supplementary Table S1

Table S1. Prediction performance of the alternative machine learning models

| Model | $R^2$ | | | MAE | | |
|---|---|---|---|---|---|---|
| | HC train | HC test | SZ | HC train | HC test | SZ |
| random forest | 0.916 | 0.895 | 0.814 | 4.460 | 3.649 | 6.469 |
| GBM | 0.983 | 0.230 | 0.312 | 1.351 | 10.382 | 11.283 |
| lasso | 0.823 | 0.282 | 0.315 | 10.182 | 9.925 | 11.272 |

#### Supplementary Table S2

Table S2. Associations between Δage and cognition after adjusting clinical characteristics

| Variable | Working memory | | Processing speed | |
|---|---|---|---|---|
| | *β* | *p* | *β* | *p* |
| group | 0.733 | 0.728 | 28.308 | <0.001*** |

| | | | | |
|---|---|---|---|---|
| sex | -1.285 | 0.119 | 5.538 | 0.055 |
| Δage | -0.173 | 0.007** | -0.541 | 0.016* |
| group*Δage | 0.116 | 0.403 | 0.341 | 0.483 |
| chronological age | -0.069 | 0.048* | -0.339 | 0.005** |
| group*chronological age | 0.050 | 0.345 | -0.348 | 0.062 |

*$p < .05$, **$p < .01$, ***$p < .001$.

### Supplementary Results by excluding participants with schizoaffective disorder

We repeated the statistical analyses for Δage using the testing data set excluding patients with schizoaffective disorder. The new results remained similar to our main analysis results (including both SZ and schizoaffective disorder patients).

As shown in Figure S3, the linear model of brain-predicted age with chronological age in HC and SZ. The overall coefficient between brain-predicted age and chronological age in SZ and HC are 1.109 and 0.966, respectively. The coefficients of two linear model was tested using *F*-test ($F(2, 258) = 17.632$, $P < 0.0001$). In young subjects, the brain-predicted ages of SZ didn’t significantly differ with HCs. The brain-predicted age of schizophrenia patients begins to be significantly greater than HC test from 30 years old. As chronological age increases, the gap of brain-predicted age between SZ and HC are larger and larger.

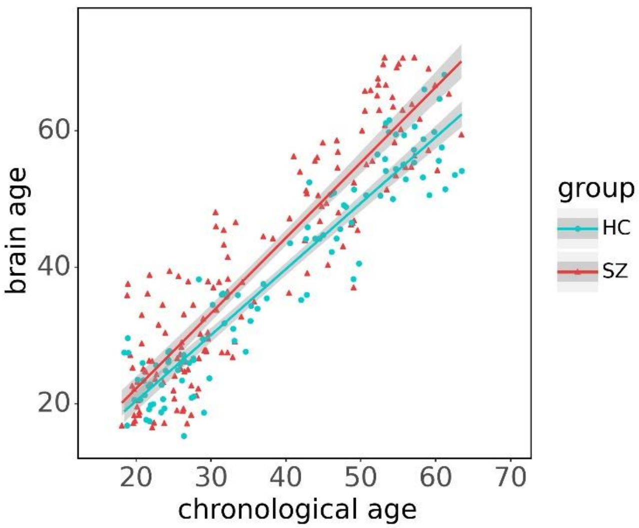


Fig. S3. Brain-predicted age and chronological age in HC and SZ with the confidence interval.

**Accelerated aging in patients**

The SZ by age interaction was positive and significant ($\beta$ = 0.143, $p$ = 0.011). The Δage was then compared between SZs and HCs for this two age groups. The Δage of the SZs was not significantly higher than that of HCs in ⩽30 age group (mean difference (95% CI): 1.890 (-0.197, 3.977), $p$ = 0.210) but was significantly higher in SZs relative to HCs in >30 age group (5.903 (3.972, 7.835), $p$ < 0.001).

**Assessing the correlation between Δage and cognition**

In patient, Δage was significantly associated with both working memory ($\beta$ = -0.168, $p$ = 0.011) and processing speed ($\beta$ = -0.485, $p$ = 0.042) after adjusting sex and chronological age (Table S3).

Table S3. Associations between three age-scaled metrics and cognition

| Group | Variable | Working memory | | Processing speed | |
|---|---|---|---|---|---|
| | | $\beta$ | $p$ | $\beta$ | $p$ |
| SZ | Sex | -1.632 | 0.127 | 7.889 | 0.043 |
| | chronological age | -0.072 | 0.044 | -0.320 | 0.014 |

| | | | | |
|---|---|---|---|---|
| Δage | -0.168 | 0.011 | -0.485 | 0.042 |
